# Label-free sensing of cells with fluorescence lifetime imaging: the quest for metabolic heterogeneity

**DOI:** 10.1101/2022.01.12.476038

**Authors:** Evgeny A. Shirshin, Marina V. Shirmanova, Alexey V. Gayer, Maria M. Lukina, Elena E. Nikonova, Boris P. Yakimov, Gleb S. Budylin, Varvara V. Dudenkova, Nadezhda I. Ignatova, Dmitry V. Komarov, Vladislav Yakovlev, Wolfgang Becker, Elena V. Zagaynova, Vladislav I. Shcheslavskiy, Marlan O. Scully

## Abstract

Molecular, morphological and physiological heterogeneity is the inherent property of cells, which governs differences in their response to external influence. The tumor cells metabolic heterogeneity is of a special interest due to its clinical relevance to the tumor progression and therapeutic outcomes. Rapid, sensitive and non-invasive assessment of metabolic heterogeneity of cells is of a great demand for biomedical sciences. Fluorescence lifetime imaging (FLIM), which is an all-optical technique is an emerging tool for sensing and quantifying cellular metabolism by measuring fluorescence decay parameters (FDPs) of endogenous fluorophores, such as NAD(P)H. To achieve the accurate discrimination between metabolically diverse cellular subpopulations, appropriate approaches to FLIM data collection and analysis are needed. In this report, the unique capability of FLIM to attain the overarching goal of discriminating metabolic heterogeneity has been demonstrated. This has been achieved using a novel approach to data analysis based on the non-parametric analysis, which revealed a much better sensitivity to the presence of metabolically distinct subpopulations as compare more traditional approaches of FLIM measurements and analysis. The new approach was further validated for imaging cultured cancer cells treated with chemotherapy. Those results pave the way for an accurate detection and quantification of cellular metabolic heterogeneity using FLIM, which will be valuable for assessing therapeutic vulnerabilities and predicting clinical outcomes.

## Introduction

Solid tumors are complex systems characterized by spatial heterogeneity at the genetic, molecular and cellular levels. Tumor heterogeneity is the critical phenomenon determining the difference in therapeutic outcomes for patients with similar histological diagnosis (1, 2). The development of omics technologies allowed for a detailed investigation of molecular pathways responsible for tumor heterogeneity, that is now considered to be a prerequisite for fast tumor growth rather than just a consequence of the neoplastic transformation and multiple mutations. Metabolic heterogeneity, i.e. the difference in cancer cells metabolism within a tumor and between tumors, is considered to be a negative prognostic factor and is accompanied by an increased probability of recurrence and higher mortality (3). The basis of metabolic heterogeneity is the ability of cancer cells to adapt to non-uniform microenvironment, e.g. local hypoxia and nutrient limitation (2), and some factors intrinsic to the cancer cells, e.g. differentiation state, proliferative activity, genetic alterations (4).

Cancer cells are capable of switching between different metabolic pathways (e.g. aerobic glycolysis and oxidative phosphorylation) depending on the local conditions. This phenomenon is known as metabolic plasticity (5). Therefore, the metabolic status may be highly variable for the cells within a single tumor and between different tumors of the same type (6–9). Evidently, visualization and quantification of metabolic heterogeneity could be helpful for optimization of cancer treatment. Hence, the methods allowing for rapid, sensitive, and non-invasive assessment of cellular metabolic heterogeneity are of a high demand for oncology.

During the last decade, fluorescence lifetime imaging (FLIM) (10), which is an all-optical technique, has proven to be a useful tool to characterize cellular metabolic state on a label-free basis. Metabolic imaging by FLIM is based on measuring fluorescence decay parameters (FDPs) of endogenous fluorophores, such as reduced nicotinamide adenine dinucleotide (phosphate) NAD(P)H and oxidized flavin adenine dinucleotide, FAD, which are involved in a number of redox reactions within the cell. It is now established that FLIM is sensitive to alterations in energy metabolism accompanying carcinogenesis (11) and cancer cells response to therapies, and that it correlates with standard biochemical and molecular assays (12–14). It was shown that FLIM enables visualization of cellular metabolic heterogeneity of cancer, both intrinsic and induced by anti-cancer therapy (15–22). However, a detailed performance comparison of different FLIM data analysis methods in quantification of metabolic heterogeneity (variability) in populations of cells has not been considered.

The FLIM-based assessment of metabolic heterogeneity at the cellular level can be reduced to the problem of discrimination between several subpopulations of cells, which differ in their FDPs. Therefore, it is important to find a way that provides the highest sensitivity to discriminate between metabolically different subpopulations of cells. Firstly, the FLIM image can be analyzed as a whole, and, consequently, the overall distribution of FDPs is analyzed over all pixels of the image. Another option is the segmentation of the image into individual objects (single cells, organelles etc.), calculation of FDPs for each object and analysis of the FDPs distribution for the segmented objects. Both approaches are used in the literature, while there is a general understanding that the latter one is more sensitive to the presence of cells subpopulations different in their metabolism (21–23). Secondly, the FLIM data can be processed either parametrically, i.e. by fitting the decay curves to a model (e.g. biexponential decay in the case of NAD(P)H and FAD), or non-parametrically. The latter approach includes the analysis of phasor plots (24, 25) or using different clustering algorithms (26), Bayesian frameworks (27, 28) and deep neural networks (29). Nonparametric methods of FLIM data analysis are extensively used due to the simplicity of interpretation of the results and the lack of necessity in data fitting, which may require calculation time. The workflow for the assessment of cancer cells metabolic heterogeneity using clustering of FDPs is schematically presented in Figure 1.

**Figure 1.**
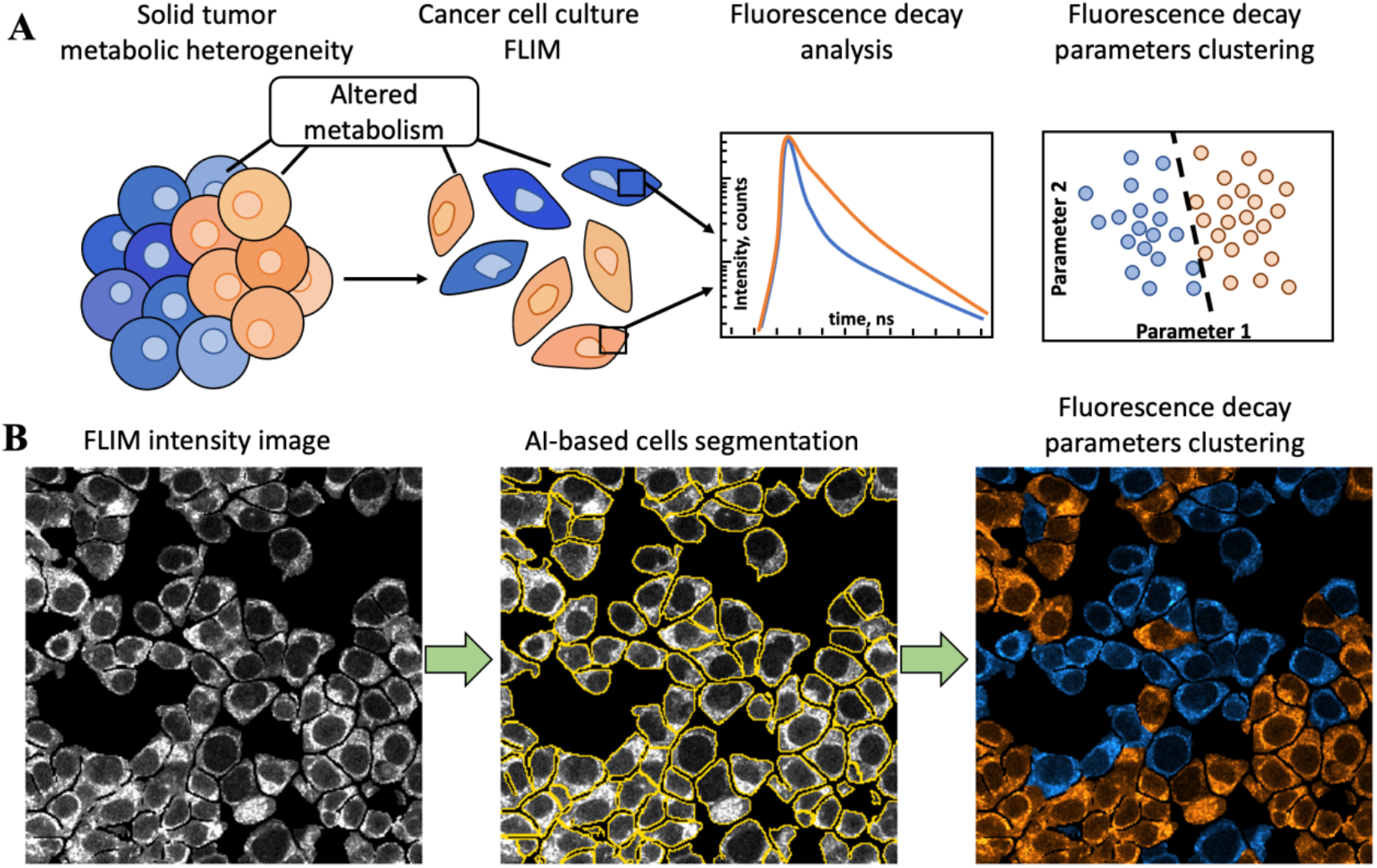
Illustration of tumor metabolic heterogeneity evaluation using FLIM. A) Cancer cells are examined using metabolic FLIM, which provides the kinetics of fluorescence decay from each pixel of the image. The obtained fluorescence decay signal depends on the metabolic state of the cell and can further be analyzed using various parametric and non-parametric methods, which can accurately predict metabolically distinct subpopulations. B) Automatic segmentation of cells in FLIM images using artificial intelligence (AI)-based approaches allows for the assessment of metabolic heterogeneity on a single cell level.

In this report, we present the results of assessment of cancer cells metabolic heterogeneity on the basis of FLIM of NAD(P)H for the simulated and experimental data using parametric (fitting to biexponential decay model) and non-parametric (phasor plot and K-means clustering) methods. By calculating the dimensionless bimodality index (BI), which characterizes the presence of two clusters of cells, we aimed at a quantitative comparison between the sensitivity of different approaches to FLIM data analysis in search of metabolic heterogeneity. To support the numerical simulation results, the metabolic heterogeneity was assessed in colorectal cancer cell line and in primary cell cultures derived from patients’ colorectal tumor upon chemotherapy. The obtained results pave the way for a more precise detection and quantification of cellular metabolic heterogeneity using FLIM.

## Results

### Analysis of bimodality in the FLIM data: numerical simulation

To assess the performance of different FLIM data analysis methods for characterization of metabolic heterogeneity, we considered the simplest yet realistic model: the system containing two clusters of cells, each characterized by a different set of FDPs. The presence of two subpopulations of cells should lead to bimodality of FDPs distribution – for instance, in the mean fluorescence lifetime distribution two peaks should be observed. However, broadening of distributions caused by dispersion of FDPs may result in the absence of visible bimodality.

The width of the distribution of the fluorescence lifetime (or other FDPs) is determined by three factors:

1. intracellular dispersion *σ_intra_*, which originates from the heterogeneity of the cell’s structure and non-uniform fluorescence lifetime distribution within the cell,
2. intercellular dispersion *σ_inter_*, which originates from heterogeneous distribution of fluorescence lifetime over individual cells within the system,
3. dispersion *σ_fit_*, which originates from the error of fluorescence decay approximation.

These three dispersions contribute to the overall width of the FDPs distribution obtained either for the whole image (i.e. distribution over binned pixels) or segmented cells (i.e. distribution over individual cells). However, the impact of *σ_intra_*, *σ_inter_* and *σ_fit_* on the width of the fluorescence lifetime distribution is different for the whole image and segmented cells analysis. Intracellular dispersion, *σ_intra_*, is lower in the latter case: each cell is characterized by a fluorescence decay curve obtained by averaging over all pixels within this cell, that reduces the dispersion of fluorescence decay parameters caused by cellular heterogeneity. The fitting error *σ_fit_* depends on the number of photons under the fluorescence decay curve N roughly as 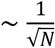 hence, it is lower when the signal is averaged over the whole cell. Intercellular dispersion of fluorescence decay parameters *σ_inter_* does not depend on the method of analysis. Hence, it can be suggested that the fluorescence lifetime distribution would be narrower in the case of the analysis for individual cells, thus resulting in a better discrimination between the subpopulations.

To verify this hypothesis, numerical simulation of the FLIM data was performed as described in the Materials and Methods section. Intercellular and intracellular dispersions were taken as *σ_intra_* = *σ_inter_* ≈ 100 ps, corresponding to typical experimentally determined parameters for NAD(P)H fluorescence (see Table S1 and the references within), and the width of the instrument response function (IRF) was set to 100 ps. Representative simulated fluorescence decay curves are shown in Fig. S1.

The numerical simulations were performed for different ratios of cells number in two clusters 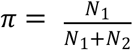 (also referred as to “Cluster 1, %”), where N_1_ and N_2_ are the numbers of cells in subpopulations, and for different values of *Δτ_mean_*, which is the distance between the median mean fluorescence lifetimes in subpopulations. The value of *π* varied in the 50÷90% range, and *Δτ_mean_* varied from 50 to 400 ps. As a measure of bimodality, the bimodality index (BI) (30) and its modifications for different FLIM processing methods were used (see the Materials and Methods section for details). The threshold corresponding to the presence of bimodality in the distribution was set to BI > 1.1 (30), while the statistical significance of the obtained BI values is calculated within the assumption that data is not bimodal, i.e. low p-value suggests the presence of the bimodality in data (the evaluation of the statistical significance of the BI is described in SI Text, Fig. S2).

We applied three approaches for FLIM data processing: fitting to biexponential decay model (Fig. 2A), phasor plot analysis (Fig. 2B) and clustering of the fluorescence decay curves (Fig. 2C). In each case, hypothesis about the presence of two modes was tested, namely, (i) distribution of the mean fluorescence lifetime was fitted to two Gaussians, (ii) the density of phasor plot population was fitted to a sum of two 2D-Gaussians and (iii) the set of fluorescence decay curves obtained for all cells was clustered into two subsets using the K-means algorithm. Next, for each of two modes the median values μ and standard deviations σ were obtained, and, based on them, the BI was calculated. The representative results of bimodality assessment using all three methods are demonstrated in Figs. 2D-F, and in Fig. S3-5 for a broad range of *Δτ_mean_* and *π*. At high *Δτ_mean_* values (~350 ps), i.e. when the difference of FDPs in two clusters of cells is large, two modes are clearly seen in the data (Fig. 2D-F, panels in a green frame). However, upon the decrease in *Δτ_mean_*, the modes corresponding to two subpopulations are superimposed, and the BI becomes low (BI < 1.1, bimodality not detected, red frame in Figs. 2D-F). Using the described approach, we performed comparison of three methods of FLIM data processing performance in the assessment of bimodality. Moreover, the analyses over whole image and segmented cells were compared.

**Figure 2.**
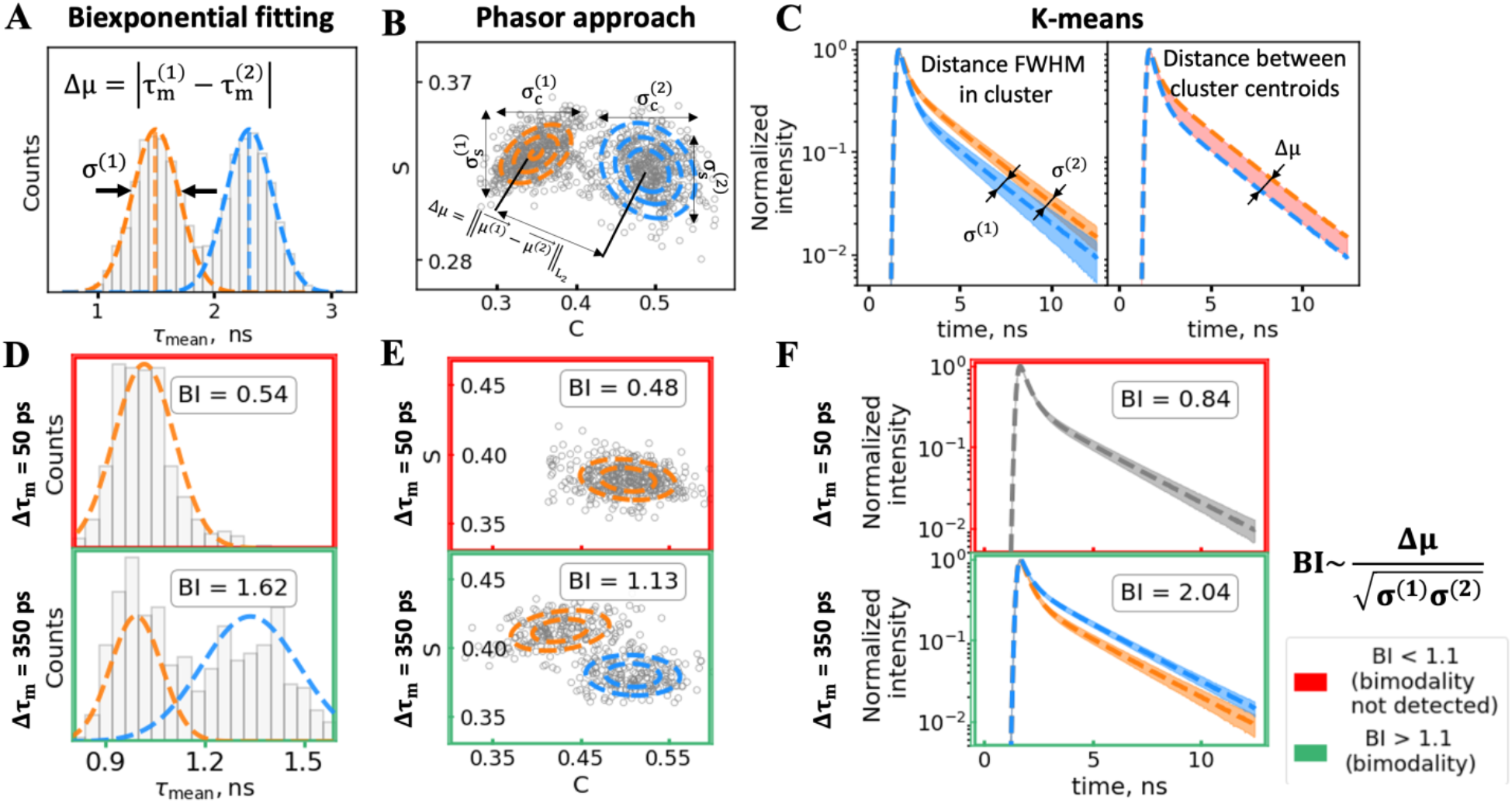
Schematic representation of bimodality assessment using A) the distribution of mean fluorescence lifetime, B) population density of phasor plot, C) K-means clustering of fluorescence decay curves. The data were obtained using numerical simulation. Under the assumption of the presence of two subpopulations, two modes are detected for each method of FLIM data analysis, and then the median value (μ) and standard deviation (σ) for each mode are used to calculate the bimodality index (BI). The results of bimodality assessment for two values of Δτ_mean_ (50 and 350 ps) using the BI estimation from D) the distribution of mean fluorescence lifetime, E) population density of phasor plot, F) K-means clustering of fluorescence decay curves. The analysis was performed over segmented cells.

As a metrics performance, we have calculated the fraction of cells Cluster 1 correctly attributed to the Cluster 1 by the algorithm: for the modeled data, this information is known *a priori.* The fraction of cells correctly assigned to Cluster 1 (with lower τmean, see Materials and Methods Section *“Clusterization performance evaluation”*) was plotted for multiple cases with different ratio of cells truly belonging to Cluster 1 π = N_1_/(N_1_ + N_2_) = 90, 70 and 50% as a function of *Δτ_mean_* (Fig. 3).

**Figure 3.**
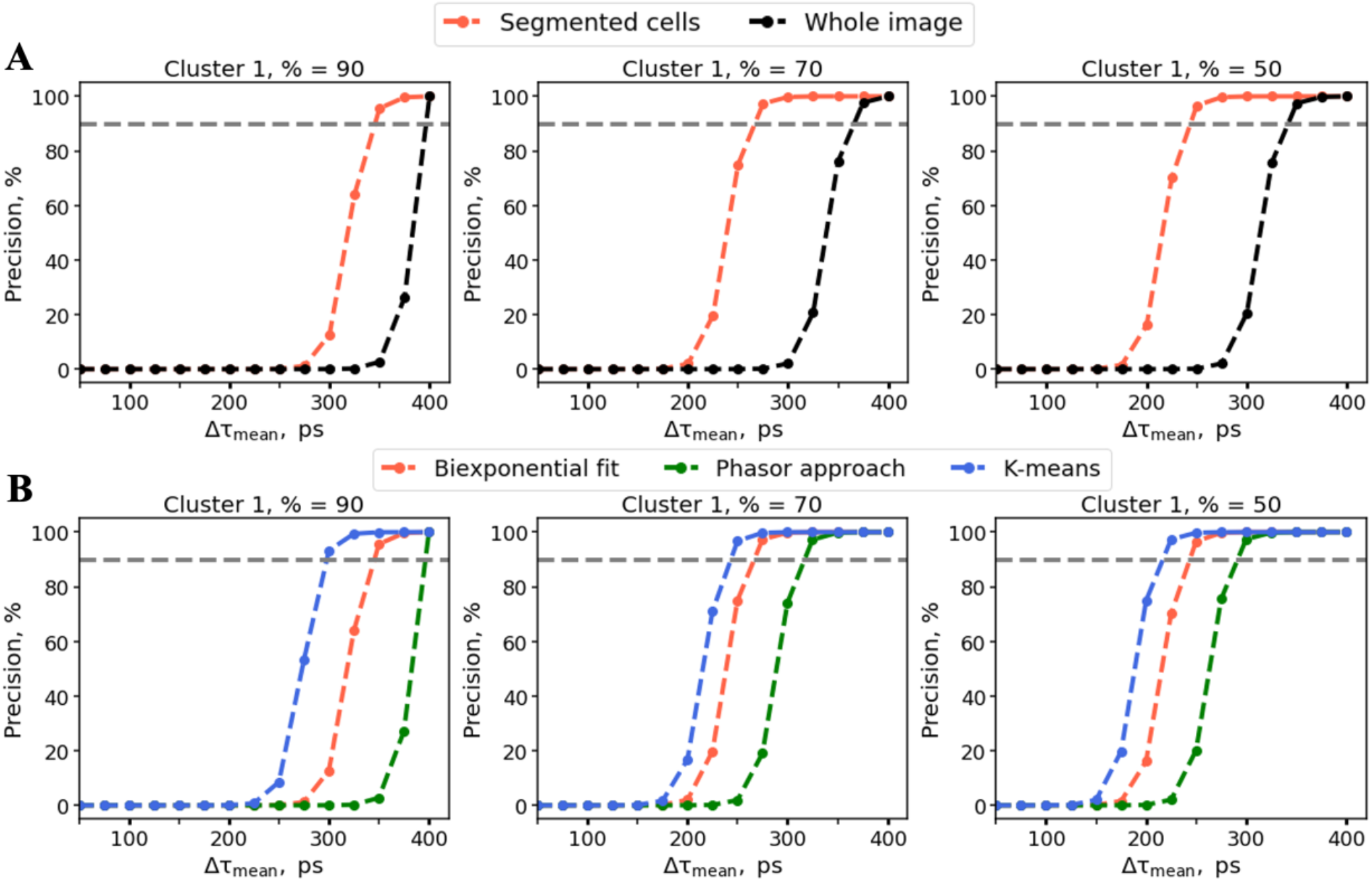
A) The dependence of the fraction of cells correctly attributed to its cluster on the distance between the median fluorescence lifetimes in the clusters (*Δτ_mean_*) obtained using biexponential fitting for whole image analysis (red) and for segmented cells (black). B) The dependence of the fraction of cells correctly attributed to its cluster on the distance between the median fluorescence lifetimes in the clusters (*Δτ_mean_*) obtained using biexponential fitting (red), phasor plot analysis (green) and K-means clustering (blue). The grey horizontal line corresponds to precision of 90% in attribution of cells to the correct cluster. The dependencies presented in panel A, B) were calculated for fraction of cells belonging to Cluster 1 equal to 90%, 70% and 50%.

Firstly, it can be clearly seen that the analysis over segmented cells always outperforms the analysis over the whole image (Fig. 3A). This observation agrees with the hypothesis that a decrease in intracellular dispersion *σ_intra_* and fitting error *σ_fit_* in the case of segmented cells analysis results in a better capability to detect the presence of two cell clusters in the system. We also note that in the case of the bin size equal to or exceeding the cell size, the fitting error would decrease, however, such binning values result in artefacts connected with mixing the signals from neighboring cells or between the cells and intercellular space. Thus, increasing the size of the bin when processing the FLIM data does not allow for better detection of cells metabolic heterogeneity (Fig. SI6).

Secondly, the results of the numerical simulation show that at high values of *Δτ_mean_* (~300 ps) all three methods of FLIM data analysis demonstrate similar performance (Fig. 3B). However, the K-means clustering exhibits the best performance compared to other methods for all of the considered fractions of the cells (Cluster 1, %) and correctly identifies the cluster to which the cell belongs at the lowest *Δτ_mean_* compared to other methods (Fig. 3B). The possibility to separate the clusters that have small differences in fluorescence lifetime parameters with high accuracy is very important when analyzing metabolic heterogeneity by NAD(P)H fluorescence lifetime (Table SI2). Our results clearly show the importance of cells segmentation for the detection of subpopulations of cells and underline the advantages of non-parametric methods, namely, K-means clustering.

### Automatic segmentation algorithm

For the further analysis of the fluorescence decay parameters of single cells on the experimentally obtained FLIM images, automatic segmentation algorithms were applied. Automatic segmentation of cells is necessary both for obtaining appropriate statistics (~1000 cells) for metabolic heterogeneity assessment in cellular populations and for creation of a rapid clinical test free of manual image analysis. To find cell borders and nuclei and segment cultured cells in FLIM images, we used the deep-learning approach based on the U-Net neural network (31) trained on the manually segmented images of the HCT116 colorectal cancer cell line with additional postprocessing of image borders to obtain single cells. Overall, the developed approach allowed us to analyse the information from 89% of single cells presented in the FLIM images of HCT116 cell lines (total amount of cells = 749) and 94% of cells in manually segmented regions of the intensity images of the patient-derived cancer cells (total amount of cells = 768). More details on the developed segmentation approach are presented in Materials and methods and SI Text.

### Analysis of bimodality in the FLIM data: experimental studies

To support the results of numerical simulations, FLIM measurements of NAD(P)H were performed for the live cultured colorectal cancer cells. In the first series of experiments, we used the human colorectal carcinoma cell line HCT116 treated for 24 hours with different doses of 5-fluorouracil, a standard chemotherapeutic agent (32).

The example in Figure 4 demonstrates that chemotherapy with 5-fluorouracil induced heterogeneous response in terms of cellular metabolism within the population of cultured cancer cells. Given the performance of different methods (Fig. 3), the analysis was performed using the mean fluorescence lifetime distribution (Fig. 4B,C) and K-means clustering of fluorescence decay curves (Fig. 4D,E) for segmented cells.

**Figure 4.**
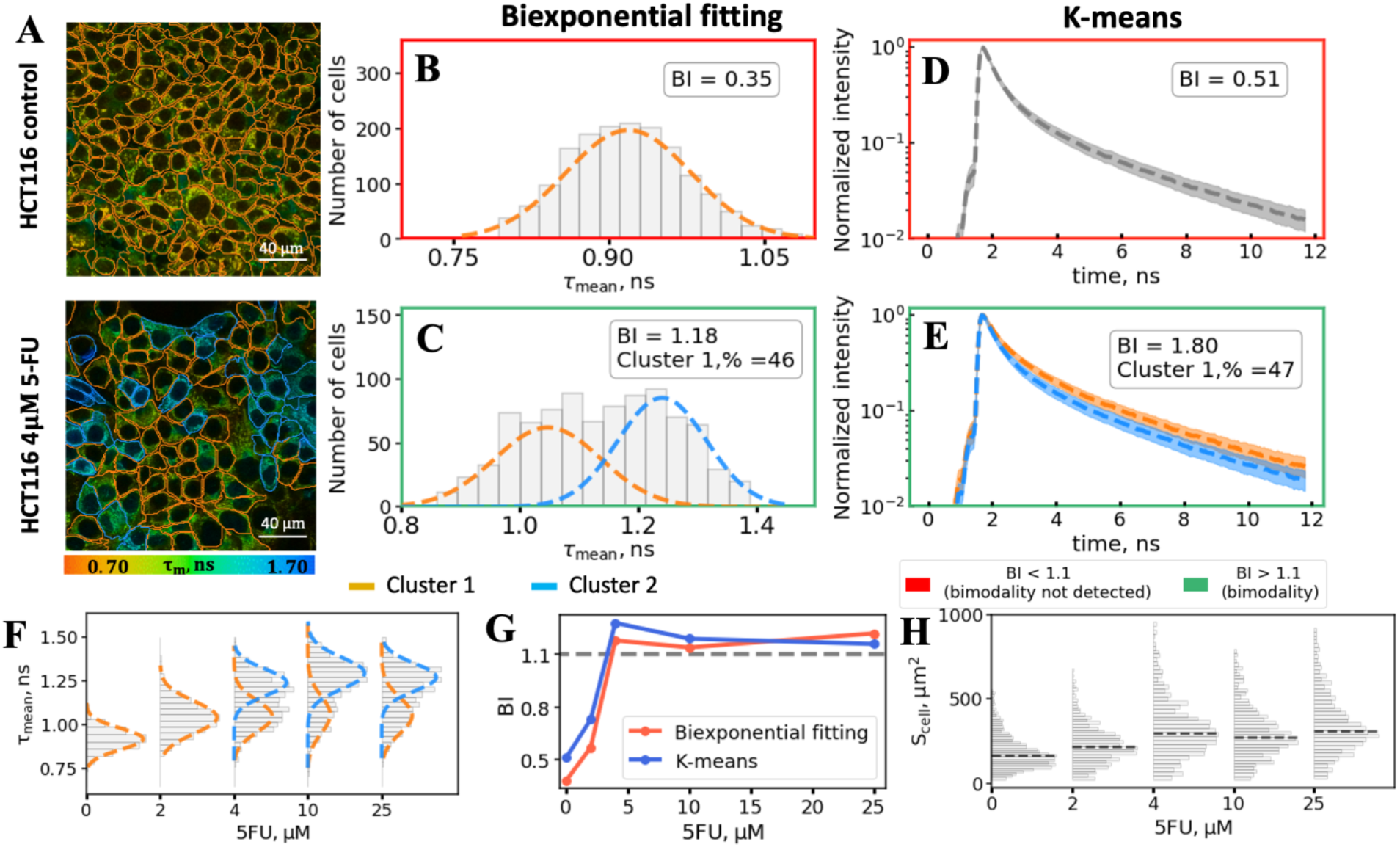
Experimental assessment of cellular heterogeneity in the human colorectal carcinoma cell line HCT116 treated with 5-FU. A) Representative FLIM images of NAD(P)H fluorescence of untreated (top panel) and treated cells (4 μM for 24 hours, low panel). Orange and blue contours correspond to two clusters as determined by the K-means algorithm applied for the fluorescence decay curves of the contoured cell. B, C) The distributions of the mean fluorescence lifetime τ_mean_ for the untreated (B) and treated (C) cells and their fits to two Gaussians. D, E) The results of K-means clustering for the untreated (D) and treated (E) samples. The two clusters of fluorescence decay curves with centroids are shown by dashed lines. F) Evolution of the mean fluorescence lifetime distribution with the drug concentration. Orange and blue contours correspond to two clusters as determined in the calculation of the bimodality index. G) The dependence of BI, calculated from fluorescence lifetime distributions (red) and using K-means clustering (blue), on the drug concentration. H) The dependence of the cells’ area on the 5-FU concentration.

It is interesting that after the treatment the distribution of the fluorescence lifetime *τ_mean_* of single cells evolved in a dose-dependent manner. At the lowest drug dose of 2 μM a small, but statistically significant shift of the mean fluorescence lifetime was observed, from 0.9 ns to 1.1 ns (t-test, p-value < 10^-4^), and the population of the cells remained metabolically homogenous (Fig. 4F). With the further increase of the dose, separation of the cells into two clusters appeared – with the small and large shifts of the mean fluorescence lifetime of NAD(P)H relative to the control, corresponding to the non-responsive and the responsive cells, respectively. Increase in the drug dose from 2 μM to 25 μM resulted in the increase of the fraction of metabolically shifted, i.e. responsive, cells up to 75%. It is important to mention that at the dose of 4 μM, close to the half-inhibitory concentration IC50 of 3 μM as determined by the MTT assay (Fig. SI7), the portion of responsive cells was ≈55%, thus confirming that the changes of the mean fluorescence lifetime of NAD(P)H are associated with the decrease of cellular metabolic activity and/or viability. Notably, the BI dependence on the drug concentration (Fig. 4G) corresponded to the viability data of the MTT assay (Fig. SI7).

Automatic segmentation of cells in the FLIM images also allowed for assessment of their morphology. Gradual increase in the cell area with 5-FU concentration was observed, without however, development of bimodality (Fig. 4H). This indicates that cellular morphology does not correlate with metabolic state of the cells and presents less reliable metric of the early response to treatment.

As a second example we analyzed metabolic heterogeneity in the cancer cell cultures obtained from the patients’ colorectal tumors. It is widely recognized that cell lines do not completely recapitulate their tumor cells of origin as inevitable selection occurs during adoption of cells to culture conditions and subsequent long-term culturing (33). From this point of view, short-term primary cell cultures represent a more ‘close-to-patient’ model.

For primary cell cultures, FLIM images of NAD(P)H were obtained within 5-7 days after isolation of cancer cells from colon tumors and tested for bimodality using three algorithms described above. The data for five patients were processed, four of which showed unimodal and one – bimodal distribution of the FDPs. Figure 5A shows the examples of primary colorectal cancer cells with uniform and heterogeneous metabolism. In the latter case, the K-means algorithm provided the highest BI value as compared to the fitting procedure and phasor plot.

**Figure 5.**
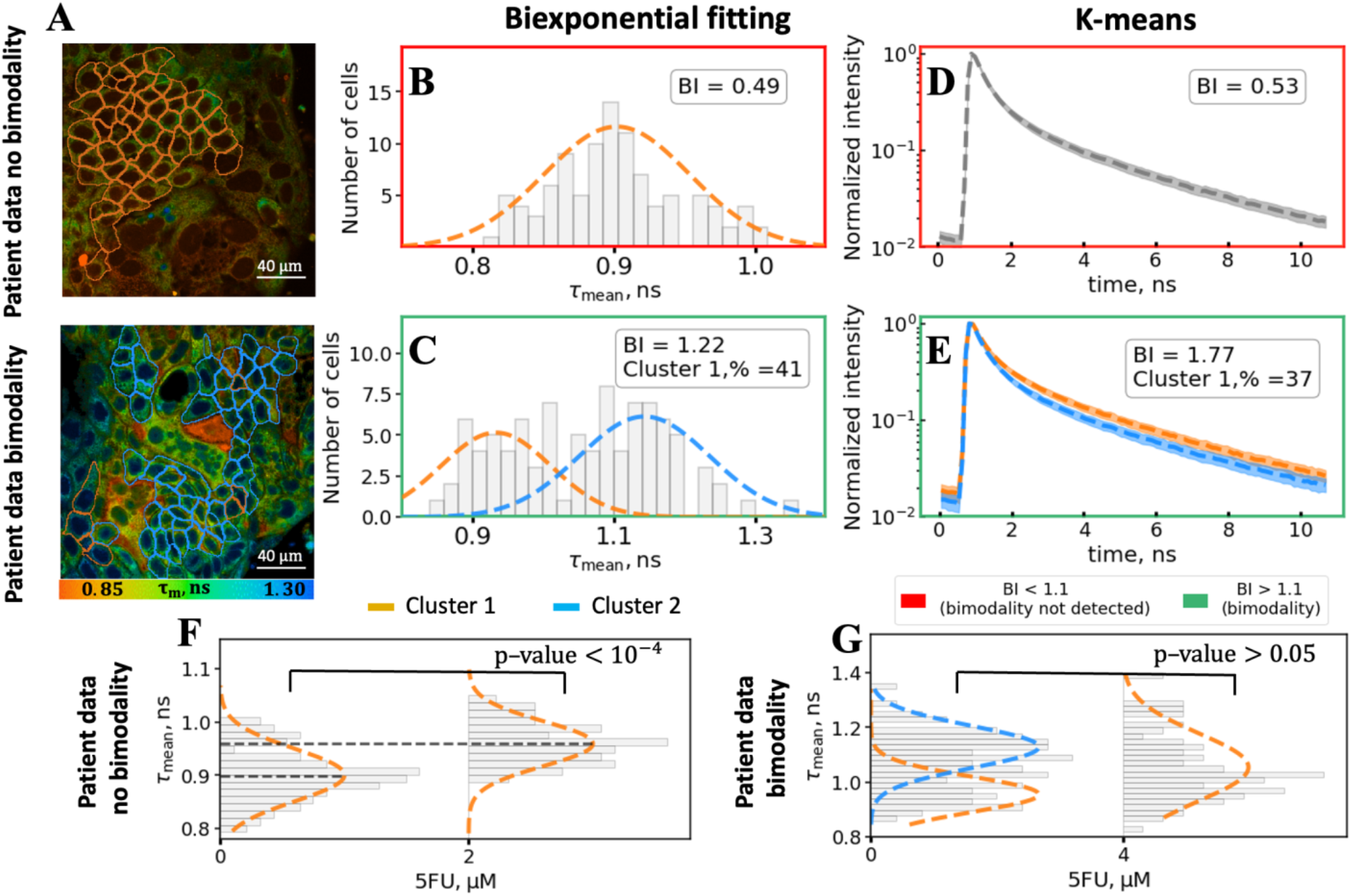
Experimental assessment of cellular heterogeneity in the primary colon cancer cell cultures. A) Representative FLIM images of NAD(P)H fluorescence of primary cell cultures exhibiting unimodal (top panel) and bimodal (low panel) metabolism. B, C) The distributions of the mean fluorescence lifetime for the primary cell cultures exhibiting unimodal (B) and bimodal (C) metabolism and its fits to two Gaussians. D, E) The results of K-means clustering for the primary cell cultures exhibiting unimodal (D) and bimodal (E) distribution of fluorescence decay parameters. The two clusters of fluorescence decay curves with centroids are shown by dashed lines. F) Changes of the mean fluorescence lifetime distribution upon treatment with 5-FU for the patient’s cells, which exhibited no bimodality. G) Changes of the mean fluorescence lifetime distribution upon treatment with 5-FU for the patient’s cells, which exhibited bimodality. Orange and blue contours in the panel A) and I) correspond to two clusters as determined by the K-means algorithm.

Further, we assessed the response of patients’ cultured cells to treatment with 2 or 4 μM of 5-FU. It was observed that for the cell cultures with initially monomodal distribution of the fluorescence lifetime (homogenous metabolism) a shift towards higher values took place (t-test, p-value < 10^−3^), similarly to what has been observed for HCT116 cell line (Fig. 5F, Table SI2). On the contrary, no statistically significant changes of fluorescence lifetime were observed for the sample, which initially exhibited metabolic heterogeneity (Fig. 5G). These results support the hypothesis that the intrinsic metabolic heterogeneity of cancer cells correlates with the poor response to therapy. In general, observation and quantification of the heterogeneity of cellular metabolism in the patient-derived material has a great clinical importance as it might be relevant for the prognosis of patients and patient-specific drug screen.

## Discussion

Imaging and quantifying tumor heterogeneity is crucially important for assessing tumor progression and tailoring cancer therapy to a specific patient. With molecular characterization and metabolic profiling, significant progress has been made in the recent years in the understanding that a high degree of spatial metabolic heterogeneity exists within tumors. Mapping and quantification of the metabolic heterogeneity at the cellular level is important not only from the point of view of fundamental aspects of carcinogenesis but also for clinical applications.

Fluorescence lifetime imaging microscopy (FLIM) of autofluorescent endogenous metabolic cofactors (NAD(P)H and flavins) allows to probe cellular metabolism by measuring their fluorescence decay time (fluorescence lifetime) in live cells and tissues non-invasively and directly observe the contrast between the different structures within the image if they have different metabolic states. Label-free principles of FLIM image acquisition and (sub)cellular resolution make this method a promising instrument to get an insight into metabolic heterogeneity of cancer cells.

Although the metabolic heterogeneity is a commonly recognized feature of tumor cells, the approaches to its quantitative assessment are poorly developed. In this study, we aimed at finding the most sensitive method to assess bimodality in the distribution of fluorescence decay parameters in cellular populations. To quantify the observed bimodality, we used the bimodality index (BI) criterion, which gives the natural measure in the case of the presence of the second mode in distribution of the estimated parameter (30). The assessment of the heterogeneity of the FLIM parameters has been previously reported using the weighted heterogeneity (wH) index (34). To the best of our knowledge, this is the only metric to quantify metabolic heterogeneity on the basis of FLIM data for today. A comparison of BI with the wH-index showed that the value of wH-index provides the results similar to BI in the heterogeneity evaluation as demonstrated in Fig. SI8 and discussed in details in the SI. Yet, the BI provides dimensionless estimation on the inherent heterogeneity of a sample, therefore it can be used to compare heterogeneity assessed by different FDPs and FLIM data analysis methods.

We note that the analysis of fluorescence intensity distribution did not reveal bimodality in the data, thus favoring FLIM as a method for the detection of metabolically distinct cells’ subpopulations (Fig. S9).

Using FLIM of NAD(P)H, we tested three approaches to the data processing for bimodality detection: (1) fitting the fluorescence decay to a biexponential decay model, (2) analysis of the FLIM data on the phasor plot, (3) clustering of the fluorescence decay curves with the K-means algorithm. Additionally, we compared the pixelwise analysis of the whole FLIM image and segmentation of the image into individual cells.

Using numerical simulation, it was demonstrated that the analysis over individual cells always outperforms the analysis over the whole image in terms of metabolic heterogeneity detection (Figs. 2–3, Figs. SI2-4). Further comparison of three above mentioned FLIM data analysis methods showed that the K-means clustering correctly identifies the cluster to which the cell belongs in the case of *Δτ_mean_* lower by as much as ~50 ps compared to biexponential fitting in the case of predominance of one of the clusters (*π* = 90%, Fig. 3). Hence, this approach occurred to be the best for detecting metabolic heterogeneity for clusters of cells with close distributions of fluorescence decay parameters.

The developed FLIM data analysis methods for metabolic heterogeneity assessment were then tested on experimental data. We showed that the extent of heterogeneity (bimodality) can be assessed using the bimodality index, and the fraction of cells in each population can be determined. In the experiments on HCT116 cancer cell line exposed to 5-FU we found that the chemotherapeutic treatment with the doses ≥4 μM induced two metabolically distinct subpopulations of cells that had been homogeneous before treatment (Fig. 4). Comparing the results of BI assessments with the MTT-assay confirmed that the metabolic changes observed by FLIM of NAD(P)H correlated with cells response to the treatment. A small shift of the mean fluorescence lifetime of NAD(P)H to longer value was assigned to non-responsive cells, while a large shift-to responsive ones. Overall, the increase of the lifetime can be a result of both inhibition of glycolysis and activation of mitochondrial respiration and presents a non-specific response to the cytotoxic drugs having different mechanisms of action (19, 35, 36). Several reports demonstrate the possibilities of FLIM to reveal heterogeneous drug response on a single-cell level. For example, in our earlier study we showed that the fluorescence lifetime of NAD(P)H is a sensitive parameter for tracking the heterogeneous cellular response to chemotherapy both in cell monolayers and in multicellular tumor spheroids (35). Specifically, metabolic heterogeneity was detected by NAD(P)H fluorescence upon treatment of HeLa (human cervical carcinoma) cells with paclitaxel – a shift to longer lifetimes developed in the more responsive cells (35). Wallrabe et al. identified heterogeneous response of prostate cancer cells AA PCa upon treatment with doxorubicin using NADH-a2%/FAD-a1% ratio (FLIRR) (37). Increased metabolic heterogeneity was shown by Shah et al. in mice with FaDu (hypopharyngeal carcinoma) tumors *in vivo* and in organoids generated from FaDu tumors after treatment with the cetuximab and cisplatin on the basis of NAD(P)H fluorescence lifetime measurements (38, 39). Although malignant cell lines are expected to be genetically and phenotypically homogeneous cell population, these studies demonstrate that chemotherapy can result in variable cellular responses. This can be associated, for example, with a variability in cell cycle phases in the “non-synchronized” cell culture or the presence of cancer stem cells, that are intrinsically more resistant to chemotherapy and heterogeneous themselves in terms of differentiation potential.

*In vitro* drug testing in cultured patient-derived tumor cells is considered as a promising approach for individualization of treatment, however there is still a lack of “ready to use” assays (40, 41). One of the issues in this methodology is the absence of reliable, quantitative metrics of cell response, which could be measured on relatively small amount of isolated cells and would take into account cellular-level heterogeneity. This motivated us to test the suggested algorithms on the short-term cell cultures generated from patient tumors. We found that, in contrast to the established cell line, primary cells can be intrinsically heterogeneous in their metabolism. For the patient-derived cells, cellular metabolic heterogeneity detected before treatment yielded in poor response to 5-FU. In general, the patient-derived cells were less sensitive to chemotherapy and displayed less pronounced metabolic changes, resembling those induced by the low dose of the drug in HCT116 cells. Previously, Sharick et al. observed cellular heterogeneity of drug response in breast cancer and pancreatic cancer patient-derived organoids. Heterogeneous metabolic response in organoids treated with the same drugs as the patient’s prescribed regimen agreed with a poor outcome for patients (42). Recently, Gillette et al. showed that a single cell metabolic heterogeneity of neuroendocrine patient-derived tumor organoids is consistent with the overall treatment response: highly heterogeneous organoids were non-responsive to all the treatments (43), which agree with our results.

The proposed approach based on the bimodality index calculation using FLIM of NAD(P)H and K-means method in combination with automatic cell segmentation has a high potential for the development of high throughput screening platforms for drug sensitivity assessments. The limitation of our approach is that only two groups of cells are considered (i.e. bimodal distribution). The introduced bimodality index could be modified to account for the presence of multiple groups at the same time. However, we did not observe more than two groups in our data, and a much larger number of cells is needed to be able to distinguish more than two groups if they are present in the population. While our investigation was focused on just one fluorescence decay parameter, τmean, the same approach can be used for other parameters like the relative contributions of free or bound NAD(P)H, or fluorescence lifetime of the bound form of NAD(P)H. In general, the underlying reasons for metabolic variability, both initial and developed after treatment, have yet to be clarified, and this will be in the focus of our future research. Further investigations of our group will also include more samples and a long-term follow-up of the patients and will be aimed to find the associations between the baseline heterogeneity of patient-derived cancer cells and treatment outcome. We also note (see SI materials and Fig. S10) that we are developing more advanced spectroscopic imaging methods based on quantum optics, which can potentially improve the accuracy of FLIM measurements while maintaining their viability of cell culture under study. In summary, the results obtained in this work provide a methodological basis for a sensitive and quantitative analysis of bimodality of fluorescence decay parameters of cells and pave the way for a more precise detection and quantification of cellular metabolic heterogeneity using FLIM. Nowadays, the choice of chemotherapy protocol for patients is determined on the basis of histological diagnosis, while individual properties of a patient’s tumor are not taken into account, which limits the efficacy of treatment. This fact stimulates the search of the early predictors of chemotherapy effectiveness. We believe that identification and accurate assessment of cellular metabolic heterogeneity of patients’ tumors with the use of suggested algorithms can become a valuable approach to develop improved, individualized treatment regimens.

## Acknowledgments

Development of the algorithms for FLIM data processing was supported by the Russian Science Foundation (project 20-65-46018). The work with primary cell cultures was supported by the Ministry of Health of the Russian Federation as part of the Government assignment (PRMU, M.V. Shirmanova PI). The authors are thankful to Irina Druzhkova, Liubov Shimolina, Vadim Elagin and Alena Gavrina (PRMU) for their kind assistance with biological studies and Prof. Vladimir E. Zagaynov (VDMC) for support of the clinical research. V.V.Y. acknowledges partial support of the National Science Foundation (NSF; DBI-1455671, ECCS-1509268, CMMI-1826078), the Air Force Office of Scientific Research (AFOSR; FA9550-15-1-0517, FA9550-20-1-0366, FA9550-20-1-0367), Army Medical Research Grant (W81XWH2010777), the National Institutes of Health (NIH; 1R01GM127696-01, 1R21GM142107-01), the Cancer Prevention and Research Institute of Texas (CPRIT; RP180588). M.O.S. acknowledges support from the Air Force Office of Scientific Research (AFOSR; FA9550-20-1-0366, FA9550-20-1-0367), the Office of Naval Research (ONR; N00014-20-1-2184), the Robert A. Welch Foundation (Grant No. A-1261), and the National Science Foundation (NSF; PHY-2013771).

The work of E.A.S., A.V.G., B.P.Y. and G.S.B. was performed under the Development program of the Interdisciplinary Scientific and Educational School of Lomonosov Moscow State University “Photonic and Quantum technologies. Digital medicine”. The work of A.V.G. on the data analysis was supported by the Foundation for the Advancement of Theoretical Physics and Mathematics “Basis” scholarship for Master students (scholarship No. 20-2-9-2-1).

## Materials and methods

### FLIM data modelling

Each cell was simulated as a square of 45×45 with a nucleus of 20×20 pixels on the image of 512×512 pixels. After that, a fluorescence decay curve was generated for each pixel based on the biexponential decay law, and the decay parameters *a*_1_ *a*_2_, *τ*_1_, *τ*_2_ were all normally distributed with relative deviation of 15% from the average within each cell. Then, the decay curves were convolved with the instrument response function (IRF) modeled as a Gaussian function with σ_IRF_ = 100 ps. After that, a Poissonian noise was added to each decay curve. Two clusters of cells were modeled by setting two sets of mean fluorescence decay parameters for each cluster.

The values of photophysical parameters of fluorescence decay curves were selected to match the experimentally observed data of NAD(P)H fluorescence lifetime imaging. The corresponding parameters were selected for the different cell compartments. The FDPs of cell cytoplasm setted as follows: 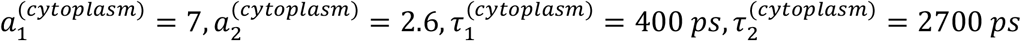; the typical nuclei FDPs: 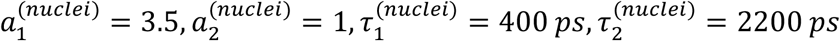, and for the background – 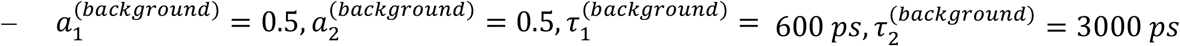. Taking the number of time bins as 500 (each corresponding to 40 ps), these parameters correspond to the average of 100 photons over the fluorescence decay (and 10 photons in maximum) for the cell, 35 photons over the fluorescence decay (and 5 photons in maximum) for the nuclei region, and 15 photons over the fluorescence decay (1 photon in maximum) for the background per pixel. The representative fluorescence decay curves from cell cytoplasm, nuclei and background are presented in Fig SI1.

To model the heterogeneity in subpopulation, values of cytoplasm FDPs of 50, 70 or 90% of cells were adjusted using the *a*_1_/*a*_2_ ratio in the way that specific *Δτ_m_* = *τ_m_*^(2)^ – *τ_m_*^(1)^ was achieved, while the number of photons under the fluorescence decay curves in average was fixed to 100. Exemplary parameters set for each subsample are presented in Table SI3.

After the modeling of the FDPs distribution, 50 cells were placed on the image of 512×512 pixels, and for each numerical experiment 10 images were considered. Two types of analysis were performed to obtain distribution of fluorescence decay parameters. In the first case, fluorescence decay parameters were obtained by fitting the decay curves in binned pixels (binning = 3, corresponding to the window of 9×9 pixels). In the second case, fluorescence decay curves were averaged for all the pixels inside each cell. The obtained fluorescence decay curves were analysed using biexponential fitting with respect to gaussian instrument response function, phasor approach and K-means clustering procedure. The amplitudes (*a*_1_, *a*_2_) and characteristic lifetimes (*τ*_1_,*τ*_2_) estimated using biexponential fitting procedure were used to calculate the mean fluorescence lifetime as *τ_m_* = (*τ*_1_ · *a*_1_ + *τ*_2_ · *a*_2_)/(*a*_1_ + *a*_2_).

### Bimodality index calculation for average fluorescence lifetime

The bimodality index (BI) for distributions of average fluorescence lifetime was calculated as follows. The each obtained *τ_m_* sample was fitted using an expectation maximization algorithm that fit the distribution of *ρ*(*τ_m_*) to the weighted mixture of two normal distributions *N*(*τ_m_*,*σ*) with different centers *τ_m_*^(1)^ and *τ_m_*^(2)^ and different standard deviations σ^(1)^, σ^(2)^, i.e.:

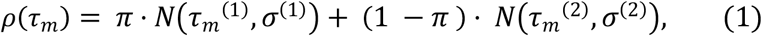

After that the bimodality index BI was calculated as (2):

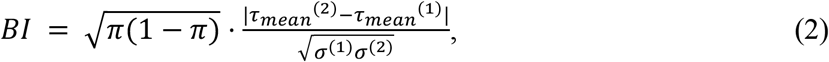

where 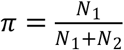.

### Bimodality index calculation for phasor parameters

The phasor analysis of fluorescence decay curves was conducted according to (24). Only the time bins with non-zero data were used to calculate C and S phasor parameters. No additional IRF corrections were applied to the data before the phasor analysis.

Similarly to the calculation of BI for distribution of average fluorescence lifetimes, the bimodality index was assessed in a two-step manner. First, the distributions of phasor parameters were fitted using expectation maximization algorithm to mixture of two normal distributions (3):

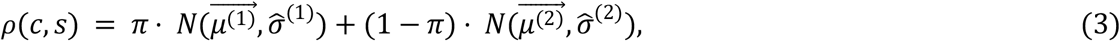

where 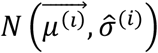 - density function of bivariative normal distribution.

After obtaining the values of centers of two Gaussians (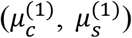 and 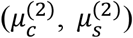) and their standard deviations (σ^(1)^ and σ^(2)^), the BI was calculated as

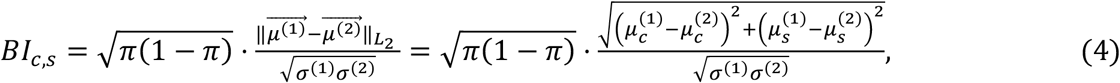

where 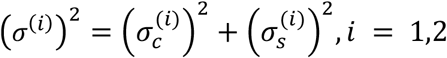.

### K-means clustering of fluorescence decay curves

The fluorescence decay curves obtained for a given numerical experiment (i.e. for fixed *π* and *Δτ_m_* values) were preliminary normalized to the [0,1] range to avoid the dependency of clustering results on the fluorescence intensity. Then, each normalized fluorescence decay curve was exponentially transformed to reduce the difference between intensities in different time bins. After that each fluorescence decay curve was treated as object with a feature-vector {*I*(*t_1_*),*I*(*t_2_*),…, *I*(*t_500_*)}, where *I*(*t_i_*) corresponds to the transformed intensity in the time bin *t_i_*. After that the K-means clustering algorithm with 2 clusters was applied to the object-feature matrix composed of preprocessed fluorescence decay curves. Using the obtained cluster labels for each decay curve and cluster centroids, i.e. fluorescence decay curves averaged within each cluster, the bimodality index was calculated as described further.

### Bimodality index calculation for clustered fluorescence decay curves

After the K-means clustering, two sets of fluorescence decay curves were obtained, and for each cluster the centroids fluorescence decay curve (centroid) was calculated. Calculation of intracluster deviations (σ^(1)^ and σ^(2)^) was performed as follows (5):

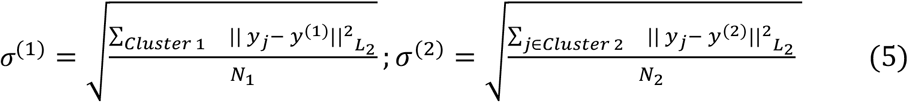

where the summation is performed for all fluorescence decay curves *y_j_* attributed either to the first or to the second cluster with centroids *y*^(1)^, *y*^(2)^, and corresponding numbers of fluorescence decay curves *N*_2_, and *N*_2_. After that, bimodality index was calculated using the *L_2_* distance between centroids *y*^(1)^, *y*^(2)^ and estimated standard deviations σ^(1)^, σ^(2)^

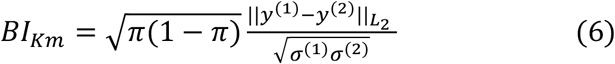

where *π* was calculated using the fraction of fluorescence decay curves attributed to the cluster with index “1”, i.e. 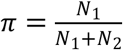.

All simulation and data analysis was performed using custom-build Python 3.7 scripts with the use of Numpy, Scipy, Scikit-Learn Matplotlib, Pandas and LmFit modules. The implementations of K-means and expectation maximization algorithms from the Scikit-learn module were used in the calculations (44).

### Cluster performance evaluation

To characterize the predictive capabilities of three methods in the assessment of cellular heterogeneity, the metrics called precision was computed as follows (7):

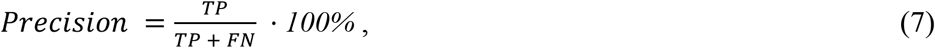

where TP – the number of cells correctly attributed to Cluster 1 (true positive), FN – the number of cells from Cluster 1 (with lower fluorescence lifetime) which were incorrectly referred to Cluster 2 (false negative).

### Cells segmentation algorithm

In this work, we implemented automatic segmentation of the human colon adenocarcinoma line HCT116 cells and cancer cell cultures obtained from the patients’ colorectal tumours. We used hybrid approach that utilizes the combination of the deep learning routines, namely, the U-Net neural network, which is widely used for binary segmentation of medical images (31, 45, 46), combined with computer vision algorithms and heuristics for object segmentation. The implemented algorithm is schematically shown in Fig. 6.

**Figure 6.**
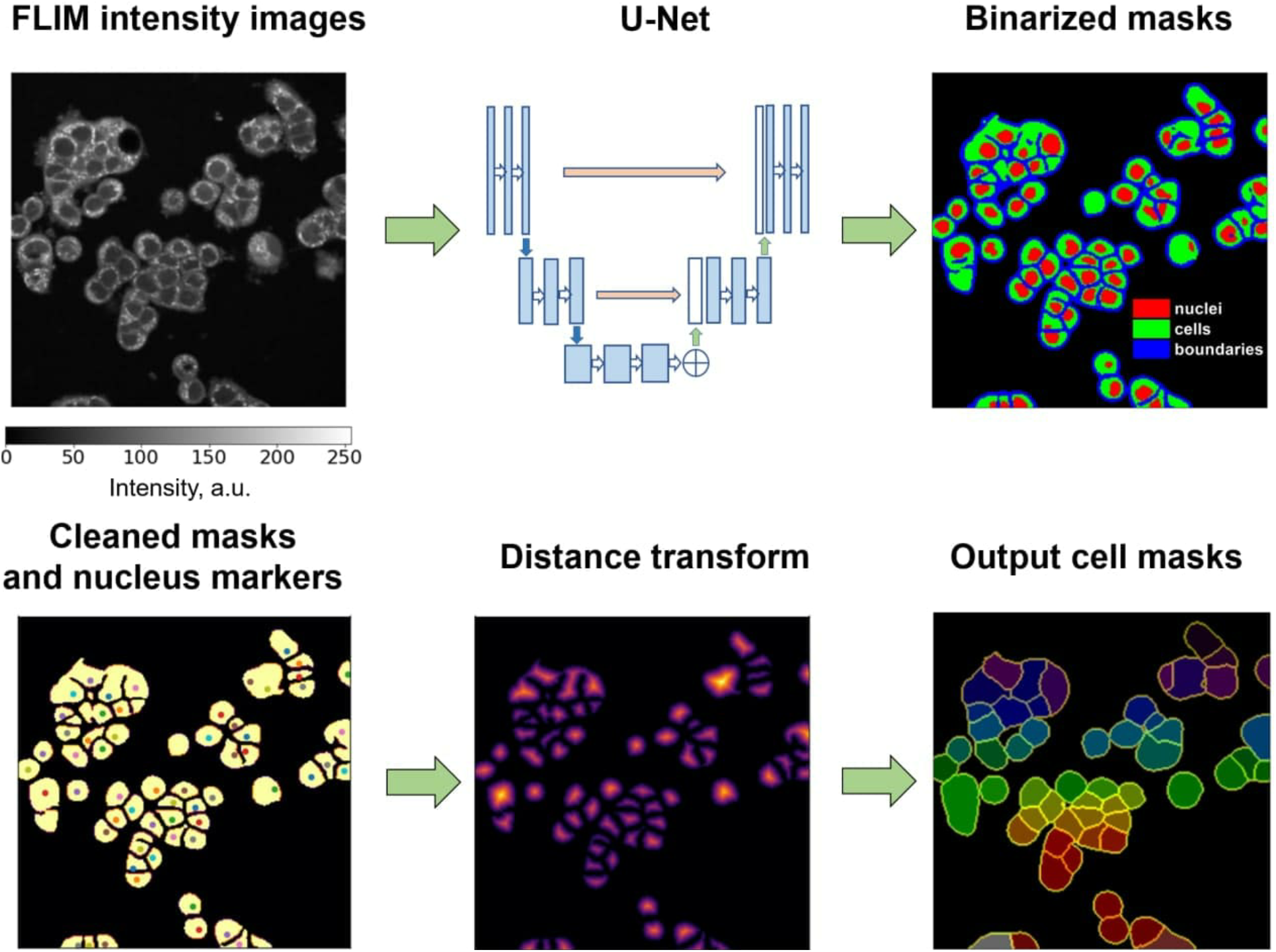
Schematic of the U-Net model-based algorithm for cells segmentation in the FLIM data.

Firstly, the intensity images were normalized to the [-0.5, 0.5] range to match the inputs of the U-net neural network. Then the pre-processed images served as an input of the multiple U-nets, which predicted cells’ borders, inner regions of cells and cells’ nuclei, i.e. each U-net for a given image predicted a map of probabilities for pixels to belong to cell’s border, cells inner region (cytoplasm or nuclei) or to cell’s nucleus. The U-nets were preliminary trained on the manually segmented images for correct probability masks prediction. The obtained probability masks were binarized and post-processed using morphological operations, and the probability masks of cells’ nuclei were used to compute the nuclei centroids. The processed binary masks and positions of the nuclei centroids were used to calculate distance maps from the nuclei centers to the cells’ borders. The obtained distance maps were further subject to the watershed algorithm, which returned the final masks.

As for the patient-derived materials exhibited artifacts related to irregular cells’ morphology and uneven sectioning, only the cells inside manually specified region were segmented, while the images of the HCT116 cells were processed as is. The details on the U-nets training, morphological operations, thresholding and other procedures are given in the SI.

### FLIM measurements

FLIM measurements on live cultured cells were performed using an LSM 880 (Carl Zeiss, Germany) laser-scanning microscope equipped with an FLIM module Simple Tau 152 TCSPC (Becker & Hickl GmbH, Germany). Two-photon fluorescence of NAD(P)H was excited with a femtosecond Ti:Sa laser (repetition rate 80 MHz, pulse duration 140 fs) at wavelength of 750 nm and registered in the range 455–500 nm. A water immersion objective C-Apochromat 40×/1.2 NA W Korr was used for image acquisition. The average power applied to the samples was ~6 mW. Image collection time was 60 s. During image acquisition, the cells were maintained at 37 °C and 5% CO_2_.

### Cell cultures

Human colorectal carcinoma cell line HCT116 was routinely grown in DMEM (PanEko, Russia) supplemented with 10% fetal bovine serum (FBS) (HyClone, USA), 2 mM glutamine, 10 mg/mL penicillin, and 10 mg/mL streptomycin. The cells were incubated at 37 °C, 5% CO2, and 80% relative humidity and passaged three times a week.

For fluorescence microscopy the cells were seeded (1×10^5^ in 2 mL) in glass-bottomed 35 mm FluoroDishes and incubated 48 hours (37 °C, 5% CO2). Then the cells were washed with PBS and placed in DMEM life medium without phenol red. The monolayer cells were incubated with 5-fluorouracil (Veropharm, Russia) at the concentrations 2, 4, 10 and 25 μM for 24 hours. Untreated cells served as a control. In total, 5 FLIM images were obtained both for the control and treated samples from randomly selected fields of view.

Surgical samples of patients’ colon tumors were provided by the Volga District Medical Center (Nizhny Novgorod, Russia) in accordance with protocols approved by the local ethical committee. All the patients included in the study had a histological verification of colorectal cancer and had not been pretreated with radiotherapy or chemotherapy. Primary cell cultures were isolated from tumors using the protocol described in ref. (47). After isolation, the cells were seeded in 96-well black/clear plates (Falcon®, Corning Incorporated, Germany) precoated with collagen I (Gibco), ~5-10 x 10^3^ cells in 200 μL DMEM per well. The medium was changed after 3 days. FLIM was performed in 5-7 days after seeding, upon cells attachment, which was controlled with light microscopy. In total, 10 FLIM images were acquired for each sample. Clinicopathological data of cancer patients are present in Table SI4.

## Authors contributions

E.A.S., M.V.S., and V.I.S. designed the study, interpreted the results, wrote the manuscript; A.V.G, E.E.N., B.P.Y., G.S.B performed numerical simulations and analyzed the data; M.M.L. and V.V.D. conducted the experiments; D.V.K. collected patients samples; W.B., V.V.Y and M.O.S. directed and supervised the study.

## Notes

### Competing Interest Statement

The authors have declared no competing interest.

