## Supplementary Information for "Label-free sensing of cells with fluorescence lifetime imaging: the quest for metabolic heterogeneity"

PNAS
www.pnas.org

### Supplementary Information for

Label-free sensing of cells with fluorescence lifetime imaging: the quest for metabolic heterogeneity

Evgeny A. Shirshin[1], Marina V. Shirmanova[2], Alexey V. Gayer[1,3], Maria M. Lukina[2], Elena E. Nikonova[1], Boris P. Yakimov[1,3], Gleb S. Budylin[1,3], Varvara V. Dudenkova[2], Nadezhda I. Ignatova[2], Dmitry V. Komarov[4], Vladislav Yakovlev[5*], Wolfgang Becker[6], Elena V. Zagaynova[2], Vladislav I. Shcheslavskiy[2,6], and Marlan O. Scully[5,7*]

Paste corresponding author name here


**This PDF file includes:**

Supplementary text
Figures S1 to S10
Tables S1 to S5
SI References

#### Supplementary Information Text

##### Determination of the statistical significance of the BI

Statistical significance (p-value) of the observed bimodality evaluated using bimodality index (BI) was calculated using bootstrap approach (1) within the assumption that the investigated parameter (e.g. $\tau_{\text{mean}}$) was unimodally and normally distributed (the null hypothesis). The alternative hypothesis suggested that the investigated parameter did not have unimodal and normal distribution. Thus, the small p-value (e.g. p-value < 0.05 or p-value < $10^{-3}$) for the observed BI could be treated as case of highly improbable unimodally distributed data, so the null hypothesis should be rejected and data should be treated as bimodal.
The empirical distribution (cumulative distribution function and probability density function) of the BI within the null hypothesis was obtained by generating random subsamples with volume N=100-500 (with step 50) $10^4$ times. The one-sided p-values of the observed bimodality index $\text{BI}_{\text{Test}}$ were calculated using cumulative distribution function of the BI by the definition, i.e.

$$p\text{-}value = Pr\,Pr\,\left(BI \geq BI_{Test} \mid H_0, N\right),$$

where the value of N was close to that in the experiment. The Fig. S2A shows the empirically derived cumulative distribution functions of the BI calculated for the subsamples with N = 250, 350, 500 and examples of the distributions of the $\tau_{\text{mean}}$ with statistically insignificant BI value (Fig. S2B), and $\tau_{\text{mean}}$ distribution that demonstrates the presence of the bimodality (Fig. S2,C).

##### Comparison of the Bimodality index (BI) and the heterogeneity index (wH-index)

To compare between the performance of the BI and the weighted heterogeneity index (wH-index), previously introduced in (6) for the assessment of metabolic heterogeneity from the FLIM data, we performed the following procedure.
Bimodal distributions of the fluorescence lifetimes were generated using the following parameters:

$$\left\{\boldsymbol{\sigma_1 = \sigma_2 = 100\,ps;\ \tau_{mean}^{(1)} = 1000ps; \tau_{mean}^{(2)} = \tau_{mean}^{(1)} + \Delta\tau_{mean};\ \ \Delta\tau_{mean} \in [100, 400]\,ps,\quad \Delta\tau_{mean}\,step = 10\,ps\ N_1 + N_2 = N = 1000; N_2 = \pi \cdot N;\ \ \pi \in [0.1, 0.5]\,\pi\,step = 0.1}\right.$$

where N is the size of the sample. For each distribution, characterized by its $\Delta\tau_{mean}$ and $\pi$ values, the BI and wH indexes were obtained, and their correlation is shown in Fig. S8. It can be seen that these two values are linearly correlated with the $R^2$ = 0.975.
We also assessed the performance of these metrics of the experimental data, obtained for the cell treated with drug. The data presented in Fig. S8,B demonstrate the full coincidence between the dependences of BI and wH-index.

##### U-net neural network training

For the prediction of the probability masks for the cells' boundaries, cells' inner regions and cells' nuclei we used three separate U-net models with architecture similar to the original work (2). To

train models we used 58 images of the HT29 and HCT116 cell lines with manually segmented binary masks of cells' borders, cells' inner region and cells' nuclei as true labels. To increase the volume of the training dataset, standard data augmentation methods were used, including image transformations: rotation, mirroring, zooming, and modulation of pixel intensity.
We used combined binary cross-entropy and dice-loss functions for the weight optimization. The training was carried out for 180 epochs until the loss function stop to decrease over 5 epochs on the training and validation datasets. At the end of training, the value of the dice coefficient was 0.85 and 0.42 for predictions of inner regions of cells and cells boundaries respectively. Relatively small value of dice coefficient for cells boundaries due to thin boundaries of 2 pixels. U-Net model used for predictions of nuclei regions trained at the same dataset images with respectively ground true during 35 epochs. At the end of training, the value of the dice coefficient was 0.56.

**Postprocessing of the U-Net model output data**

To determine the boundaries and inner regions of cells, the probability maps predicted by U-net models were thresholded using the Otsu thresholding algorithm. Further, artifacts were filled on the binarized masks of the cell areas using the morphological opening, closing and erosion and dilation algorithms. A distance transform was used to determine the centres of the cells and delimit the adjacent cells. Finally, the standard markers-based Watershed algorithm was used to obtain the individual inner regions and borders of the cells, which allows obtaining simply connected areas of individual cells. The initial coordinates of markers were chosen as centres of cells' nuclei, obtained with additional network. The processing procedure was performed using the custom-made script in Python 3.7 with the numpy, scipy, skimage and tensorflow libraries.

**Evaluation of the performance of the automatic segmentation algorithm**

The performance of the developed U-net model in cells segmentation was evaluated as follows. For the 10 images of HCT116 cells' and 10 images of the patient-derived samples from the test-set not used for the U-net model training we estimated the fraction of cells not detected by the developed automatic segmentation algorithm. After that we calculated the fraction of cells that was not detected as single cells, i.e. multiple cells were merged in the one region and was attributed to the one label in segmentation mask. The results of the performance are presented in Table S5.
We note that the performance of the used automatic segmentation algorithm for the patient-derived samples was evaluated only for the regions of the images that were manually segmented by ImageJ software.

**Quantum-enhanced fluorescence lifetime measurements**

One problem in biophotonics is the damage to the living sample due to intense laser light. Two-photon absorption with entangled light provides a thought-provoking approach to this problem. This can be understood in a simple form by considering the two-photon fluorescence as in Fig. S10. The interaction Hamiltonian for the process of Fig. S10 is

$$H = \hbar g a_3^\dagger a_2 a_1 + adj, \qquad \text{(S1)}$$

where $\hbar g$ is the coupling energy, and $a_i$ $(i = 1,2,3)$ is the field annihilation operator (8). Working in the Heisenberg picture, we have

$$\dot{a}_3 = \frac{i}{\hbar}[H, a_3] = -iga_2a_1, \qquad \text{(S2)}$$

so that for weak enough coupling in a thin slab,

$$a_3(L) \cong a_3(0) - i\eta a_2(0)a_1(0), \qquad \text{(S3)}$$

where $\eta = \frac{gL}{c}$; i.e. the time of interaction $\tau = \frac{L}{c}$.
Now suppose that the photons 1 and 2 are in the two-photon squeezed state

$$|\Psi_0\rangle = \left[\frac{|0_1 0_2\rangle + \beta|1_1 1_2\rangle}{\sqrt{1+\beta^2}}\right]|0_3\rangle. \qquad \text{(S4)}$$

Then the photon number in modes 1 and 2 is given by

$\langle n_1 \rangle = \langle \Psi_0 \rangle = \frac{\beta^2}{1+\beta^2} \equiv \underline{n},$ (S5)

and $\langle n_2 \rangle$ is the same. However, if we calculate $\langle n_3 \rangle$ using Eq. (S1) and (S4) we find

$\langle \Psi_0 \rangle = \frac{\eta^2 \beta^2}{1+\beta^2} \equiv \eta^2 \underline{n},$ (S6)

which is somewhat counterintuitive because one might have thought that since $\langle \Psi_0 \rangle \cong \eta^2 \langle \Psi_0 \rangle$, which leads to $\eta^2 \langle n_1 \rangle \langle n_2 \rangle$, one might have expected $\langle n_2 \rangle \cong \eta^2 \underline{n}^2$, which is wrong. Thus, by using the entangled light described by Eq. (4) we get a much larger two photon fluorescence (going as $\underline{n}$) than we would have for a separable state like $|n_1 n_2\rangle = |n_1\rangle \otimes |n_2\rangle$, which would give a two-photon fluorescence going as $\underline{n}^2$, when $\langle n_1 \rangle \cong \langle n_2 \rangle = \underline{n}$. The point being that for $\underline{n} < 1$ and taking a high flux of photon pairs the entangled state would give a much stronger fluorescence than would a high flux of separable photon pairs since $\underline{n}^2 \ll \underline{n}$. The use of entangled light provides a much more efficient way to excite fluorescence reducing the photodamage. Once those entangled photons are employed, different detection strategies can be used (see, for example, the most recent work by Mukamel et al (9)) to extract to temporal properties of fluorescence emission leading a novel way of fluorescence lifetime imaging, which can be extended to the current studies.

**Fig. S1.** The representative fluorescence decay curves from cell cytoplasm, nuclei and background simulated for numerical experiment.

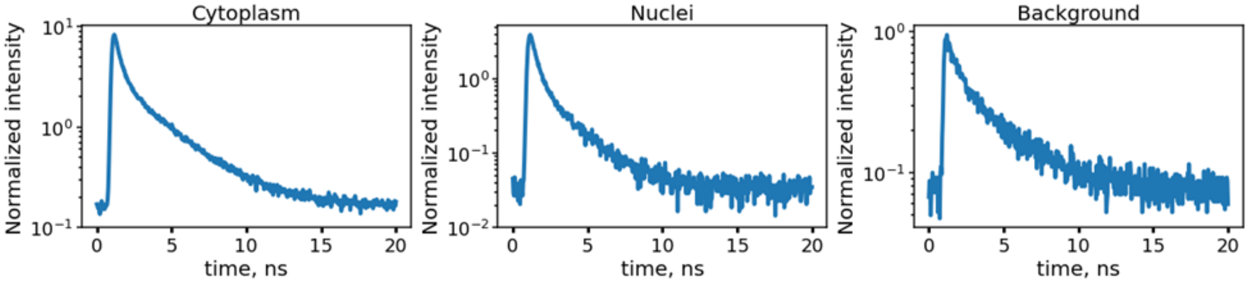

**Fig. S2.** A) Cumulative density function of the BI. The results of the p-value assessment for the obtained BI values in the cases of unimodal (B) and bimodal (C) distributions. For the bimodal

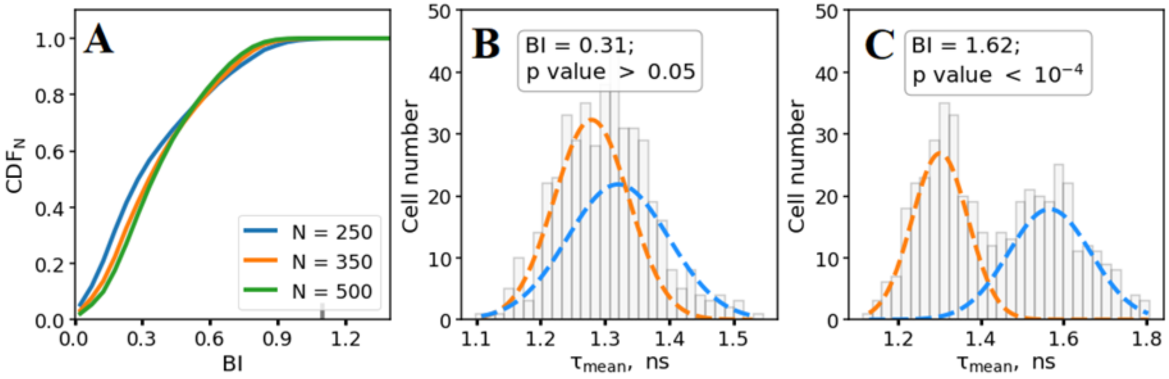

**Fig. S3.** Numerical simulation of distributions of the mean fluorescence lifetime calculated for different ratios of cells in two subpopulations (π, or "Cluster1, %") and distance between the mean fluorescence lifetimes of two clusters ($\Delta\tau_{mean}$). The distributions were calculated for the analyses using segmentation of cells (top panel) and whole image analysis (low panel). Red and green colors correspond to BI < 1.1 (no bimodality detected) and BI > 1.1 (bimodal distribution), respectively.

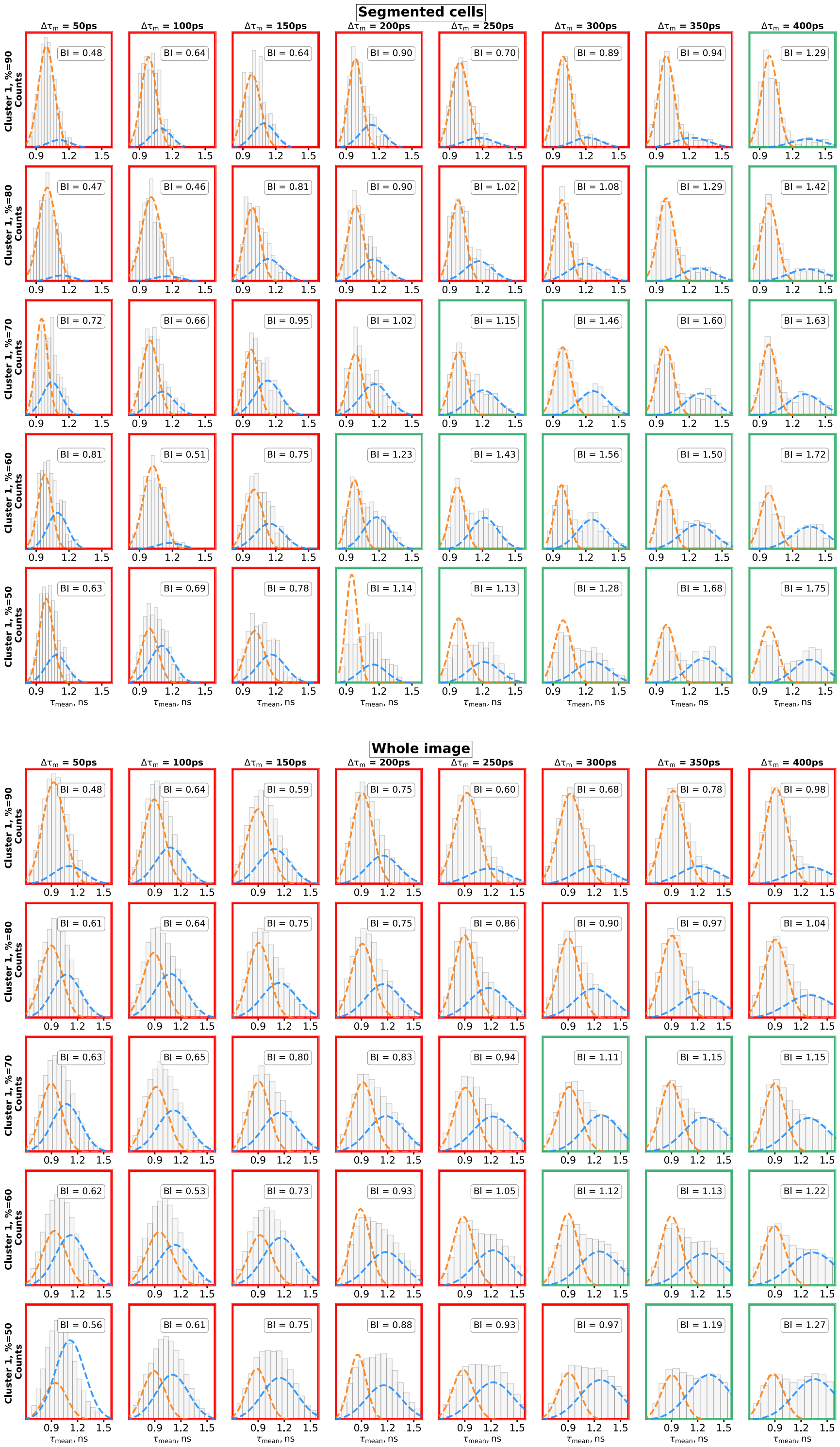

**Fig. S4.** Numerical simulation of data distributions on a phasor plot calculated for different ratios of cells in two subpopulations (π, or “Cluster1, %”) and distance between the mean fluorescence lifetimes of two clusters ($\Delta\tau_{mean}$). The distributions were calculated for the analyses using segmentation of cells (top panel) and whole image analysis (low panel). Red and green colors correspond to BI < 1.1 (no bimodality detected) and BI > 1.1 (bimodal distribution), respectively.

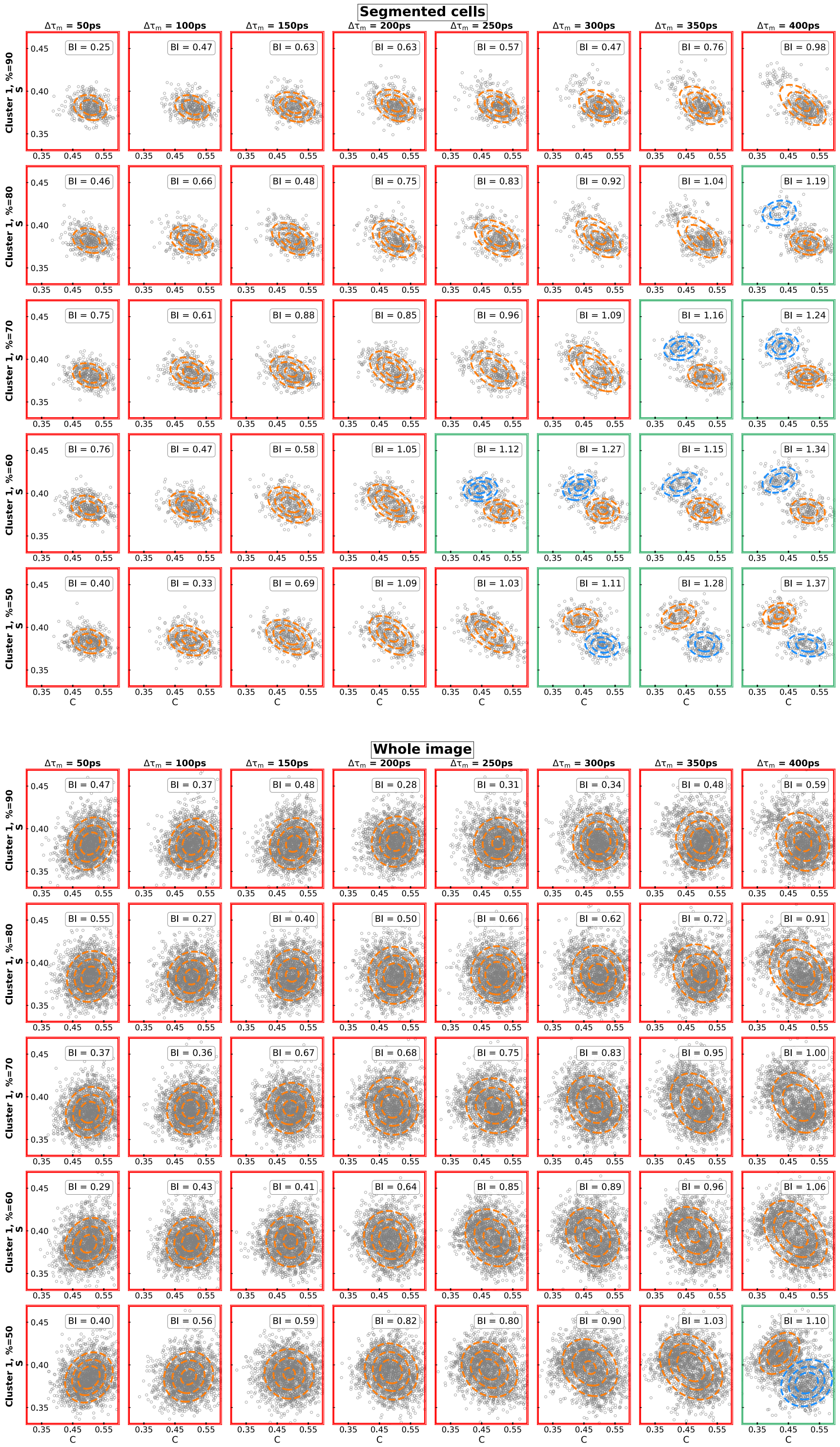

**Fig. S5.** The analysis of simulated data using K-means calculated for different ratios of cells in two subpopulations (π, or “Cluster1, %”) and distance between the mean fluorescence lifetimes of two clusters ($\Delta\tau_{mean}$). The representative fluorescence decay curves and centers of clusters (dashed lines) obtained using the K-means algorithm calculated for segmented cells.  C) The representative fluorescence decay curves and centers of clusters (dashed lines) obtained using the K-means algorithm calculated for the whole image.

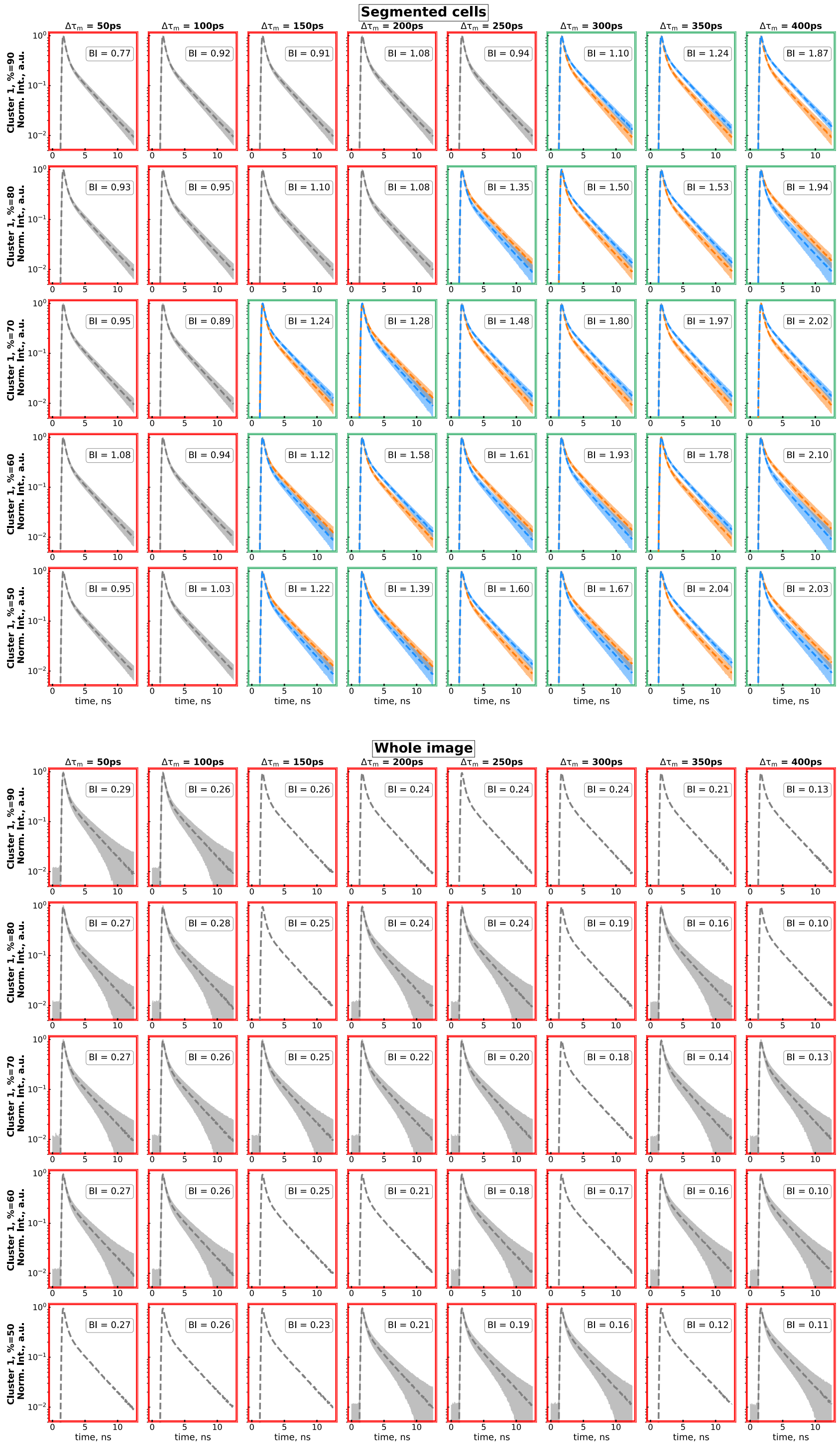

**Fig. S6.** Numerical simulation of distributions of the mean fluorescence lifetime calculated for different ratios of cells in two subpopulations (π, or "Cluster1, %") and distance between the mean fluorescence lifetimes of two clusters ($\Delta\tau_{mean}$). The distributions were calculated for the analyses using whole image analysis with binning = 14 (900 pixels) which is comparable with the typical cell size (~2000 pixels) used in the numerical experiment. Red and green colors correspond to BI < 1.1

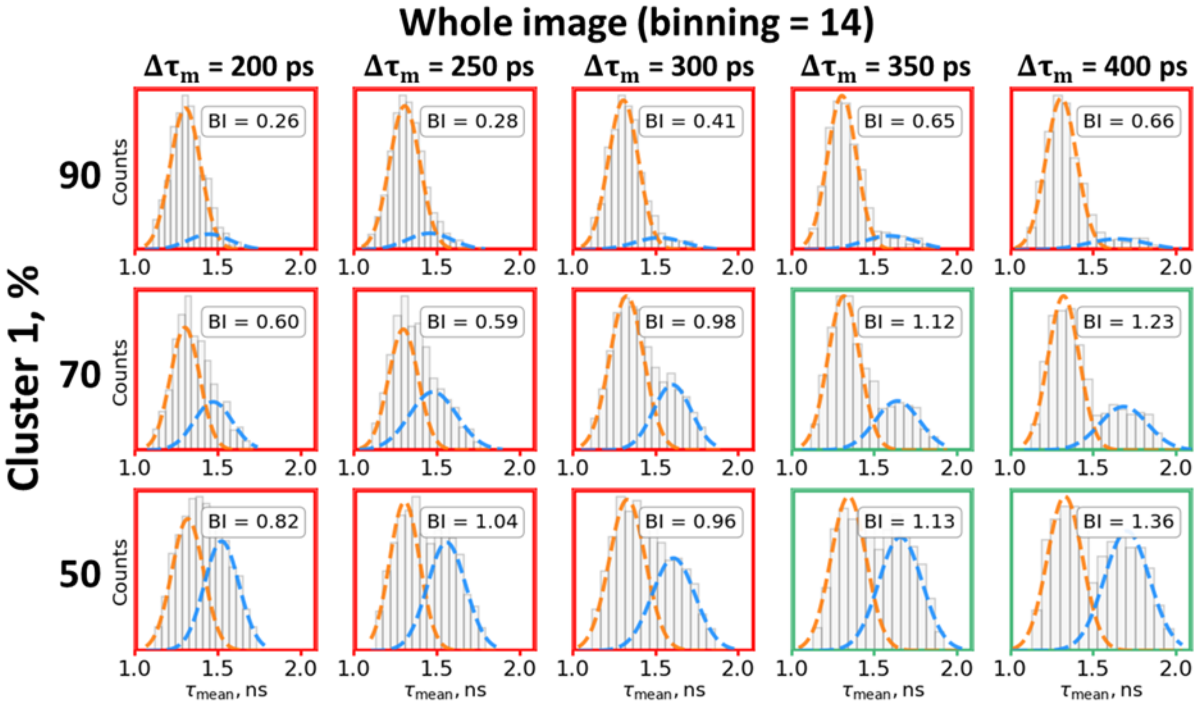

**Fig. S7.** MTT-assay of viability of HCT116 cells in the presence of 5-fluorouracil.

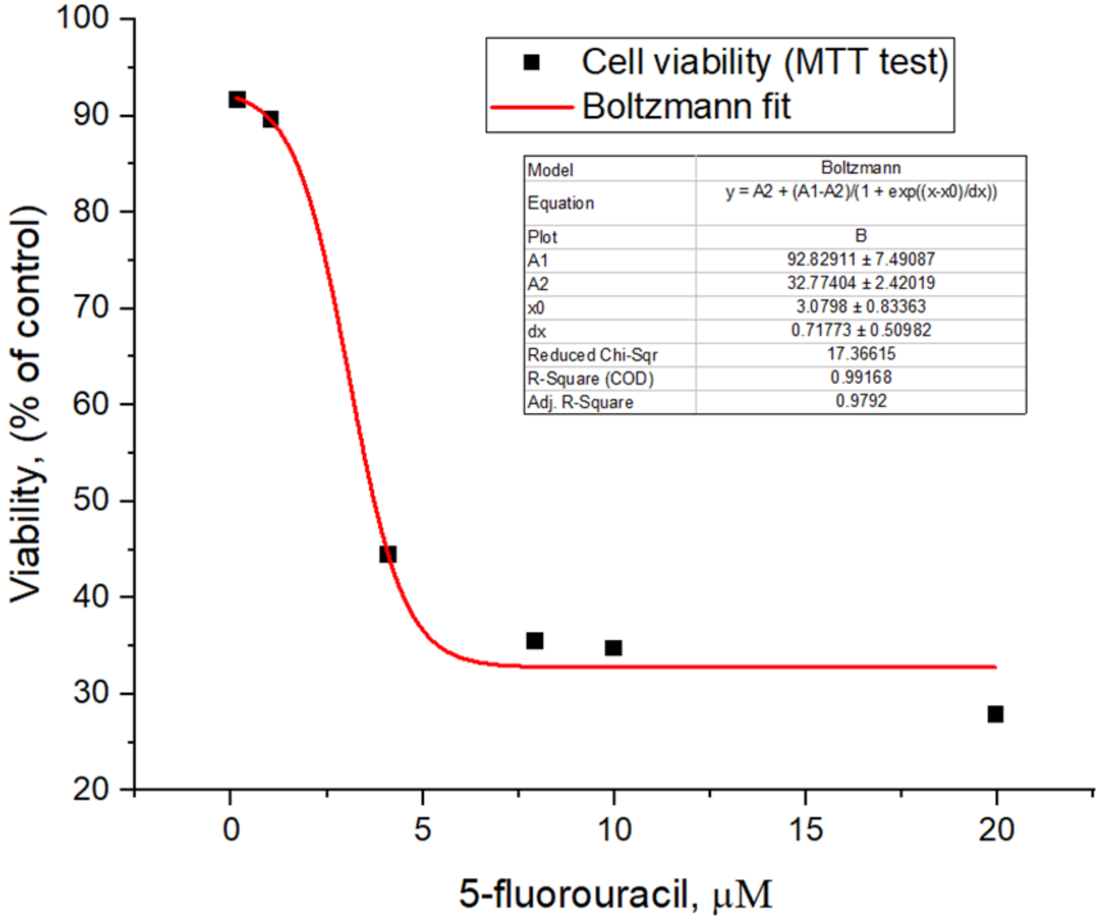

**Fig. S8.** A) Correlation between the BI and wH-index obtained for the modeled data. B) The dependences of the BI (obtained using K-means clustering) and wH-index on the concentration of

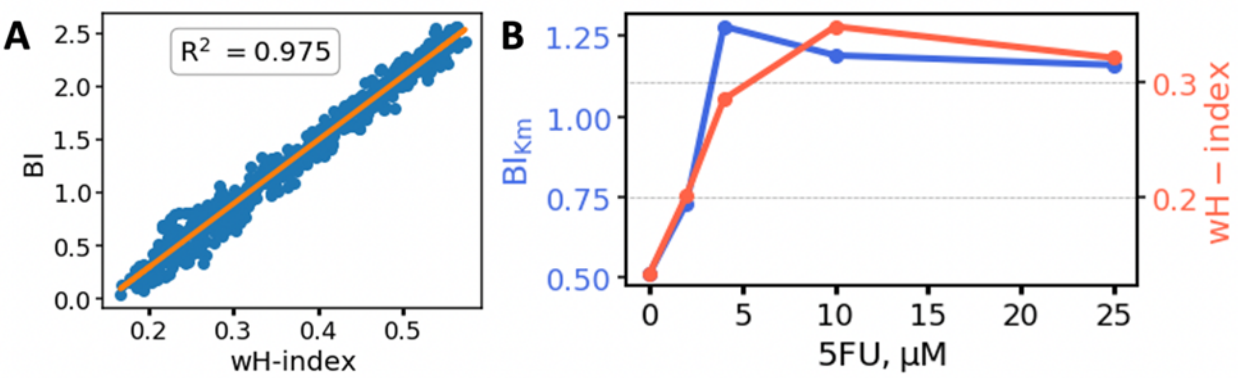

**Fig. S9.** A) The distribution of the logarithmic fluorescence intensity for the cells that exhibited heterogeneous metabolic response as assessed using fluorescence decay parameters. The log transform for the fluorescence intensity was performed to prepare data for the bimodality analysis. The bimodality index estimated for the presented distribution is equal to 0.81. B) The distribution of the fluorescence lifetime $\tau_m$, and the fit to two Gaussians of the experimental distribution. The value of the BI = 1.33 corresponds to pronounced heterogeneity in the data.

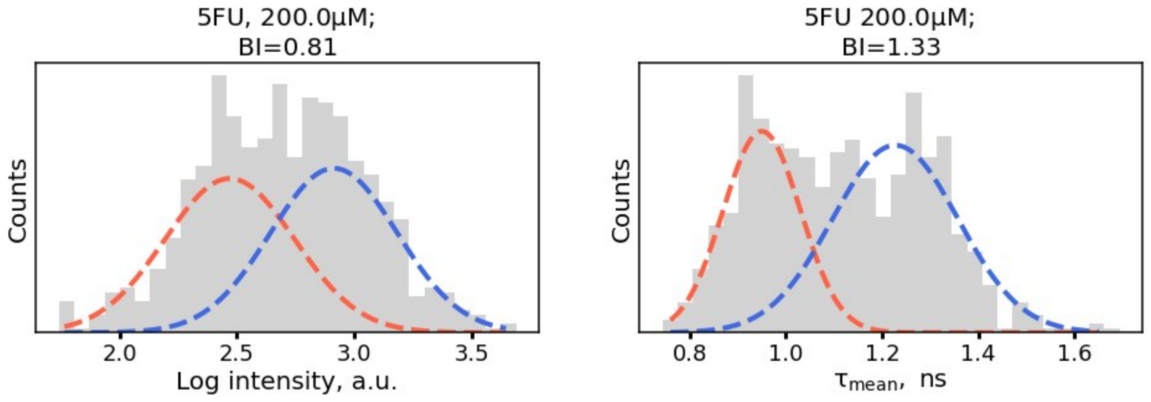

**Fig. S10.** Energy diagram illustrating the concept of quantum-enhanced fluorescence lifetime imaging. Two photons having frequencies $\nu_1$ and $\nu_2$ are resonant with the atomic transitions of frequencies $\omega_{bc}$ and $\omega_{ab}$ and the resulting fluorescence $\nu_3$ at $\omega_{ac}$.

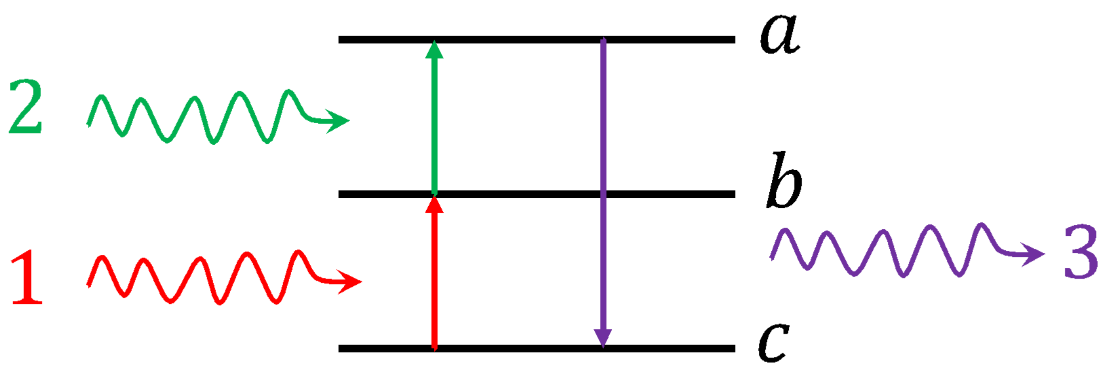

**Table S1.** Representative values of the standard deviation of the mean fluorescence lifetime distribution ($\sigma_{inter}$) and the distance between the median mean fluorescence lifetimes in metabolically heterogeneous subpopulations of cancer cells ($\Delta\tau_{mean}$) obtained in the NAD(P)H FLIM experiments.

| Experiment | $\Delta\tau_{mean}$ and $\sigma_{inter}$ | Reference |
|---|---|---|
| Two subpopulations of breast cancer cells (SKBr3 & MDA-MB-231) | $\Delta\tau_{mean}$ = 450 ps<br>$\sigma_{inter}$ ~ 100 ps | (3) |
| Cancer cells response to chemotherapy | $\Delta\tau_{mean}$ = 400 ps<br>$\sigma_{inter}$ ~ 100 ps | (4) |
| Coculture of two cell lines | $\Delta\tau_{mean}$ = 300 ps<br>$\sigma_{inter}$ ~ 100 ps | (5) |
| Cancer cells response to chemotherapy | $\Delta\tau_{mean}$ = 100 ps<br>$\sigma_{inter}$ ~ 100 ps | (6) |
| Cancer cells response to chemotherapy | $\Delta\tau_{mean}$ = 370 ps<br>$\sigma_{inter}$ ~ 140 ps | (7) |

**Table S2.** BI and NAD(P)H fluorescence lifetime assessments for the patients-derived cancer cells.

| n/n | Bimodality before treatment | Bimodality after treatment | 5-fluorouracil concentration, μM | NAD(P)H $\tau_{mean}$(ns) before treatment (mean±SD) | NAD(P)H $\tau_{mean}$(ns) after treatment (mean±SD) | P-value |
|---|---|---|---|---|---|---|
| 1 | No | No | NA | 0.99±0.02 | NA | |
| 2 | No | No | 2 | 0.95±0.07 | 0.98±0.06 | 0.0013 |
| 3 | No | No | 4 | 0.90±0.05 | 0.99±0.07 | $<10^{-4}$ |
| 4 | No | No | 2 | 0.95±0.08 | 1.01±0.08 | $<10^{-4}$ |
| 5 | Yes | No | 4 | 1.07±0.10 | 1.01±0.20 | 0.186 |

**Table S3.** Typical FDP obtained for a simulated fluorescence decay curve in a single pixel ($\tau_1$ = 400 ps, $\tau_2$ = 2700 ps).

| parameters | $\Delta\tau_{mean}$ = 50 ps | $\Delta\tau_{mean}$ = 100 ps | $\Delta\tau_{mean}$ = 200 ps | $\Delta\tau_{mean}$ = 300 ps | $\Delta\tau_{mean}$ = 400 ps |
|---|---|---|---|---|---|
| $a_1$ cytoplasm – cluster 1 | 7.4 | 7.4 | 7.4 | 7.4 | 7.4 |
| $a_2$ cytoplasm – cluster 1 | 2.6 | 2.6 | 2.6 | 2.6 | 2.6 |
| $a_1$ cytoplasm – cluster 2 | 7.17 | 6.96 | 6.52 | 6.09 | 5.65 |
| $a_2$ cytoplasm – cluster 2 | 2.83 | 3.04 | 3.48 | 3.91 | 4.35 |

**Table S4.** Clinicopathological data of cancer patients included in the study.

| n/n | Tumor Site | Tumor Type | Tumor Stage | Differentiation |
|---|---|---|---|---|
| 1 | Primary, cecum | Adenocarcinoma | T3NxM1 (stage IV) | High |
| 2 | Primary, sigmoid | Adenocarcinoma | T3N1M0 (stage III) | Moderate |
| 3 | Primary, sigmoid | Adenocarcinoma | T4N2M1 (stage IV) | Moderate |
| 4 | Primary, descending colon | Adenocarcinoma | T4N2M1 (stage IV) | Moderate |
| 5 | Primary, sigmoid | Adenocarcinoma | T2N0M0 (stage II) | Moderate |

**Table S5.** Performance of the developed U-Net model in cells segmentation determined for the validation dataset of HCT116 cells (n = 749) and patients' cells (n = 768).

| | Detected | | Not detected |
|---|---|---|---|
| | Single cells | Multiple cells merged in one region | |
| HCT116 cell line images | 90% | 4.2% | 5.8% |
| Patient-derived cancer cell images | 94% | 3% | 3% |
